# vClassifier: a toolkit for species-level classification of prokaryotic viruses

**DOI:** 10.1101/2024.05.28.596318

**Authors:** Kun Zhou, James C. Kosmopoulos, Karthik Anantharaman

## Abstract

As the most abundant and diverse biological entities, prokaryotic viruses play pivotal roles in ecological systems. Their taxonomic classification has been instrumental in elucidating their diversity and ecological functions. However, determination of viral taxonomy remains a considerable challenge. Recently developed approaches succeed in assignment of viral taxonomy at higher ranks, such as at the family level and above, but struggle at the subfamily level and below to the genus and species resolutions. Here, we describe the vClassifier toolkit, a phylogeny-informed methodology to provide species-level taxonomic assignments of viruses. We used single-copy marker genes relevant to specific taxa and reference phylogenetic trees for these groups which facilitates direct comparisons with the taxonomic framework of the International Committee on Taxonomy of Viruses (ICTV). Our method demonstrated significant congruence with the ICTV taxonomy, showing 84−91% alignment at the subfamily and genus levels. For species-level classification, our strategy was integrated with average nucleotide identity, yielding a high congruence rate of over 92% with the taxonomic data from the NCBI Virus database. Benchmarking comparisons revealed that vClassifier matches or surpasses other available tools regarding precision and classification success rates. By achieving objectivity and high levels of consistency, vClassifier streamlines the taxonomic categorization of prokaryotic viral genomes. Accurate assignments at the subfamily, genus, and species levels will significantly refine the taxonomic resolution of viruses, fostering a deeper understanding of viral diversity in microbiomes and ecosystems.

## 1 Introduction

Prokaryotic viruses, which infect bacteria and archaea, constitute the most abundant biological entities on our planet [1]. Their influence on ecosystem dynamics is profound, as they significantly shape global biogeochemical cycles and life on Earth [2]. Moreover, these viruses play a critical role in regulating microbial populations, promoting genetic diversity through horizontal gene transfer, and driving evolutionary processes [3]. By lysing host cells, they release organic matter that fuels nutrient cycles, thereby maintaining ecological balance [4].

However, despite their ecological importance, the vast diversity of prokaryotic viruses has historically been challenging to categorize [5]. Traditional methods for classifying these viruses have been limited by the lack of suitable genetic markers and the immense genetic variability found among viral populations [6]. Recent advancements in computational methodologies for deducing the taxonomy of prokaryotic viruses have revealed bacteriophages and archaeal viruses are vastly diverse. These computational techniques have predominantly utilized gene content, network-based criteria, genome-centric phylogenetic criteria, and pairwise distances and associated thresholds [7].

Prominent tools for classifying prokaryotic viruses at the family level and above include geNomad and vConTACT [8-11]. For classification at the subfamily and genus levels, tools such as vConTACT [8, 9, 12], GRAViTy [13, 14], VICTOR [15], ViPTree [16], and VirClust [17] have been developed. vConTACT generates viral protein clusters and analyzes gene-sharing networks across whole genomes. In contrast, GRAViTy leverages genome features like gene content and orientation for systematic analyses. VICTOR integrates phylogenetic and clustering approaches, utilizing BLAST to compute intergenomic distances with the Genome Blast Distance Phylogeny method [18]. Similarly, ViPTree also constructs phylogenetic trees based on pairwise genomic distances. Aside from the other tools, VirClust operates without the need for a reference and builds viral genome trees through protein clustering. These tools have proven to be sensitive and accurate in the study of prokaryotic viruses. Nonetheless, challenges remain in taxonomic classification. For instance, VICTOR tends to be slow and lacks scalability, constrained by the number of genomes it can process, and it may not fully account for the monophyly of taxa within trees. Additionally, vConTACT and GRAViTy do not sufficiently consider viral phylogenies, which are pivotal for delineating evolutionary relationships critical to taxonomy [19]. Moreover, ViPTree and VirClust do not perform bootstrapping for large datasets. Lastly, accurate species-level assignments are crucial for understanding the specific roles and impacts of individual viral species within their ecosystems [20]. However, no software is dedicated to the species-level assignment of viral genomes from human and environmental viromes metagenomes, posing a significant challenge for detailed viral taxonomy and ecological studies.

Genome-based phylogenies are ideal for capturing evolutionary relationships, making the criteria of monophyletic clusters and tree topology apt for taxonomic delineation. This approach is well-established in prokaryotes using conserved single-copy genes (SCGs), as demonstrated by tools such as GTDB-Tk [21, 22]. For dsDNA prokaryotic viruses within the *Caudoviricetes* class, SCGs are known to be conserved within certain taxonomic ranks, notably families and subfamilies, such as *Ackermannviridae* and *Ounavirinae* [23]. This conservation suggests that members within a family or subfamily possess a similar set of single-copy genes, hinting at the potential universality of SCGs in lower taxonomic ranks of viral taxa. Here, we identified single-copy markers specific to a broad spectrum of taxa, spanning from genera to realms. We constructed reference genome trees using concatenated sequences of these single-copy proteins. Our findings demonstrated that the phylogenetic topologies of taxa are highly consistent with the ICTV taxonomy at the subfamily and genus levels.

We incorporated this phylogeny-based strategy into the taxonomic assignment of prokaryotic viruses and present vClassifier, an automated and objective toolkit that places viral query genomes into concatenated protein reference trees specific to families or subfamilies. vClassifier assigns taxonomic classifications to genomes of bacteriophages and archaeal viruses based on the monophyly of viral groups within these trees. Moreover, vClassifier integrates average nucleotide identity to determine species-level classifications. When benchmarked against other programs, vClassifier demonstrated comparability or superiority in performance. Overall, we demonstrate that viruses contain single-copy marker genes that can be used to obtain automated accurate species-level assignments using vClassifier.

vClassifier is freely available for download at https://github.com/AnantharamanLab/vClassifier.

## 2 Methods

### 2.1 Taxa and marker selection

We accessed the MSL38 v1 release of the ICTV taxonomy system (April 25, 2023). Our selection criteria focused on complete genomes with corresponding GenBank accessions, which were also classified according to the ICTV taxonomy. We compiled a dataset of viral genomes with these selected GenBank accessions from the National Center for Biotechnology Information (NCBI) database comprising 5,128 genomes belonging to bacterial and archaeal viruses. These genomes spanned 212 distinct taxa and were organized into eight taxonomic ranks defined by the ICTV taxonomy. To ensure robustness in our analysis, we included only those taxa for which at least 10 genomes were available, as detailed in **Table S1**.

For identifying suitable markers within each taxon, we retrieved profile hidden Markov models (HMMs) of virus orthologous groups from the Virus Orthologous Groups database (VOGdb, release vog216) at http://vogdb.org, accessed on March 31, 2023. We then screened the protein dataset, downloaded from the database, for these profile HMMs using HMMER version 3.3.2 [24]. The screening process adhered to stringent parameters established previously: an E-value threshold of 1 × 10^−3^ and a minimum alignment coverage of 50% [23]. Ultimately, a range of 1 to 276 markers were conserved across 203 taxa, detailed in **Supplementary Tables S1** and **S2**. The selection of these markers was based on specific criteria: (1) occurrence in 50% or more of the genomes of bacterial and archaeal viruses within a taxon; (2) an average copy number not exceeding 1.2 per genome to ensure the single-copy nature; and (3) an average protein length greater than 100 amino acids, to facilitate reliable phylogenetic analysis.

### 2.2 Sequence alignment and phylogenetic inference

For each taxon, single-copy gene markers were identified using HMMER version 3.3.2, following the methodology described above. The sequence with the highest bit score was designated as the best hit. In instances of multiple hits for a marker, the sequence exhibiting the highest score was selected to ensure the highest fidelity in our phylogenetic analysis. To improve the precision of our data, each Multiple Sequence Alignment (MSA) of markers was individually refined using trimAL [25]. This refinement involved trimming columns that appeared in less than 50% of the taxa, enhancing the overall quality of the alignments.

Following this, we concatenated the refined individual alignments, inserting gaps in the sequences where markers were absent from certain genomes. This concatenated MSA was further processed to exclude genomes that represented less than 5% of the total alignment length in amino acids, ensuring that only highly representative sequences were included in our phylogenetic analysis. The final reference tree was inferred from the refined MSA using Fast-TREE version 2.1.11 [26] under the LG + GAMMA model. To provide robust support for the phylogenetic branches, we implemented 1000 nonparametric bootstrap replicates. The resultant phylogenetic trees were rooted at their midpoints [27], providing a reliable baseline for tree interpretation. In total, 203 phylogenetic trees were generated, each representing the evolutionary relationships within a specified taxon.

### 2.3 Benchmarking and comparison against ICTV classification systems

We assessed the consistency of our phylogenetic method with ICTV classification system. To facilitate this benchmarking, we accessed ICTV taxon information (MSL38 v1 release). Each ICTV taxon comprising two or more representatives was defined as a phyletic group. Moreover, an ICTV taxon was considered monophyletic if it included at least two genome representatives and achieved a bootstrap value of 75% or higher, as determined by the topology within the respective tree for each taxon. The accuracy of our method was quantified by the ratio of the number of monophyletic groups to the total number of phyletic groups within each taxon.

For the comparative analysis at the subfamily and genus levels, we utilized ViPTree [16], VirClust [17], vConTACT3 [11], which were run with default parameters to cluster viral genomes associated with 36 families or 55 subfamilies, as detailed in Table S3. The congruence with ICTV taxonomy was subsequently compared among ViPTree, VirClust, vConTACT3, and our newly developed tool, which we refer to as vClassifier. This comparative approach allowed us to evaluate the effectiveness and precision of vClassifier in aligning with established taxonomic standards.

### 2.4 Workflow of vClassifier

#### 2.4.1 Placement of genomes in reference trees

vClassifier processes genome assemblies affiliated with the 36 families or 55 subfamilies, accepting data in FASTA format. The tool uses Prodigal v2.6.3 with options (-m -p meta) to predict genes [28]. Following gene prediction, single-copy marker genes from the genome queries are identified and aligned using HMMER version 3.3.2, employing the previously described method. The sequence with the highest bit score is considered the best hit, and in instances of multiple hits, the sequence with the highest score is selected for further analysis. The identified marker genes are then aligned using MAFFT v7.505 [29]. This alignment process ensures that the sequences are accurately organized according to their evolutionary relationships. The aligned sequences are subsequently concatenated into a comprehensive multiple sequence alignment. This alignment is further refined by trimming columns represented in less than 50% of taxa using trimAL [25], ensuring that only the most conserved and relevant genetic information is retained. The final step involves placing the genomes into reference phylogenetic trees that were previously generated (see details in **Sequence alignment and phylogenetic inference)** using pplacer [30]. **Fig 1** describes the overall workflow of vClassifier.

**Fig 1.**
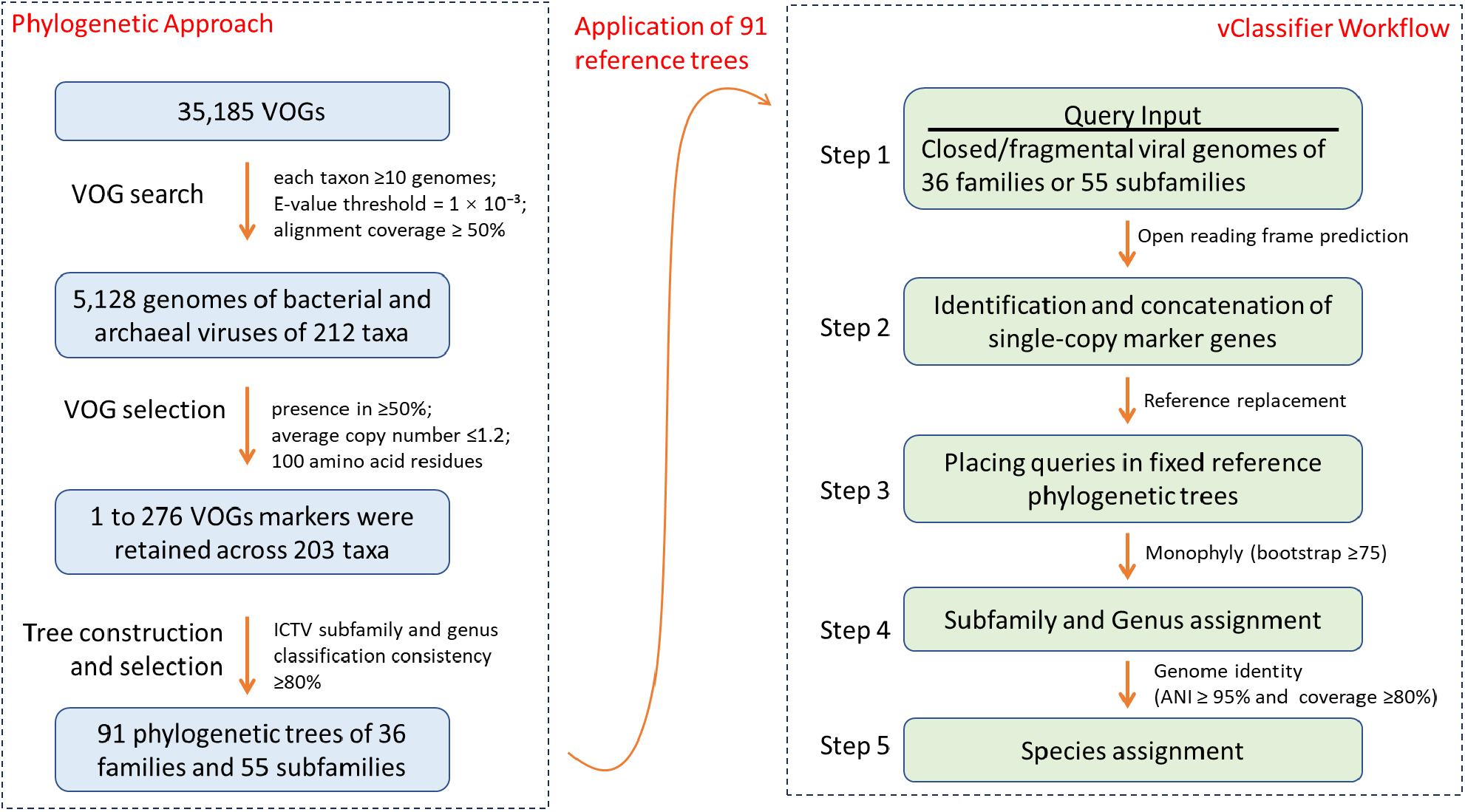
Flowchart of phylogenetic reconstruction and vClassifier workflow. The left box represents the initial phase, where phylogenetic trees are constructed based on viral SCGs derived from Viral Orthologous Groups (VOGs). Taxa: Biological classification ranks such as species, genera, families, etc.

#### 2.4.2 Subfamily/genus/species assignment

The classification of a query genome at the subfamily or genus level depends on its precise positioning within the reference phylogenetic tree. If every reference genome within a monophyletic cluster achieves a bootstrap value of 75% or higher and all belong to the same reference subfamily or genus, the query genome in the cluster is then assigned to that particular subfamily or genus. This method ensures that the classification is both accurate and reflective of established phylogenetic relationships, providing a robust basis for taxonomic assignment.

For the classification of a query genome at the species level, alignment coverage and average nucleotide identity (ANI) are critical factors. Species assignments are determined using ANI, calculated with FastANI v.1.33, applying customized parameters (--fragLen 500 –minFraction 0.8) [31]. Specifically, if a query genome is positioned within a genus, it is assigned to the species of the closest reference genome that demonstrates high alignment similarity. This determination requires that the reference genome meet stringent criteria: an alignment fraction of at least 80% and an ANI of 95% or higher. Additionally, the species assignment is based on the highest composite alignment score, calculated as the product of the alignment fraction and the ANI (fraction × ANI), ensuring the most accurate and reliable taxonomic placement.

#### 2.4.3 Requirements

vClassifier is engineered to function optimally on a server outfitted with multiple CPUs and 128 GB of RAM. It efficiently handles the classification of nearly 10,000 genomes within approximately 20 hours, utilizing 30 CPUs. For optimal results, it is recommended to use vClassifier with genomes that are of high quality; specifically, those with an estimated completeness of over 90%, as evaluated by CheckV [32]. This ensures that the genomes being classified are robust and comprehensive, leading to more accurate and reliable taxonomic assignments.

#### 2.5 Benchmarking

We evaluated vClassifier’s proficiency in classifying viruses from diverse environments, including aquatic, host-associated, sediment, and soil contexts. Utilizing a dataset of 9,724 viral scaffolds sourced from the IMG/VR database [33], we determined the taxonomic assignment rate for prokaryotic viruses across 36 families and 55 subfamilies based on the outcomes generated by vClassifier. The final output file generated by vClassifier included taxonomic information for each query viral scaffold at the species, genus, subfamily, family, order, class, phylum, kingdom, and realm levels. To statistically compare the classifications at the subfamily and genus levels, we selected vConTACT3 and employed it with its default parameters, enabling a comprehensive analysis of vClassifier’s effectiveness relative to established tools.

## 3 Results

### 3.1 Numerous single-copy gene markers in viral taxa at low ranks

Based on our search against VOGdb, we identified a total of 3,009 protein families serving as single-copy markers prevalent across the selected 203 taxa spanning five viral realms (as detailed in **Table S1** and illustrated in **Fig 2A**). Interestingly, aggregating the single-copy genes from each taxon across the five realms revealed that no markers were universally shared among all realms. However, one marker was shared between the realms *Varidnaviria* and *Duplodnaviria*, and four markers were shared between *Adnaviria* and *Duplodnaviria* (shown in **Fig 2B**). This finding underscores the absence of universal single-copy markers, akin to the prokaryotic 16S ribosomal RNA genes, across viral realms [34]. Nonetheless, identifying these single-copy marker genes across a broad spectrum of viral taxa provides a valuable resource for taxonomic and evolutionary studies [35]. This dataset highlights the diversity in marker gene distribution, reflecting the varied evolutionary histories of these viruses. Such heterogeneity in marker gene distribution has significant implications for the depth and reliability of taxonomic classifications, enhancing our understanding of viral phylogeny and taxonomy.

**Fig 2.**
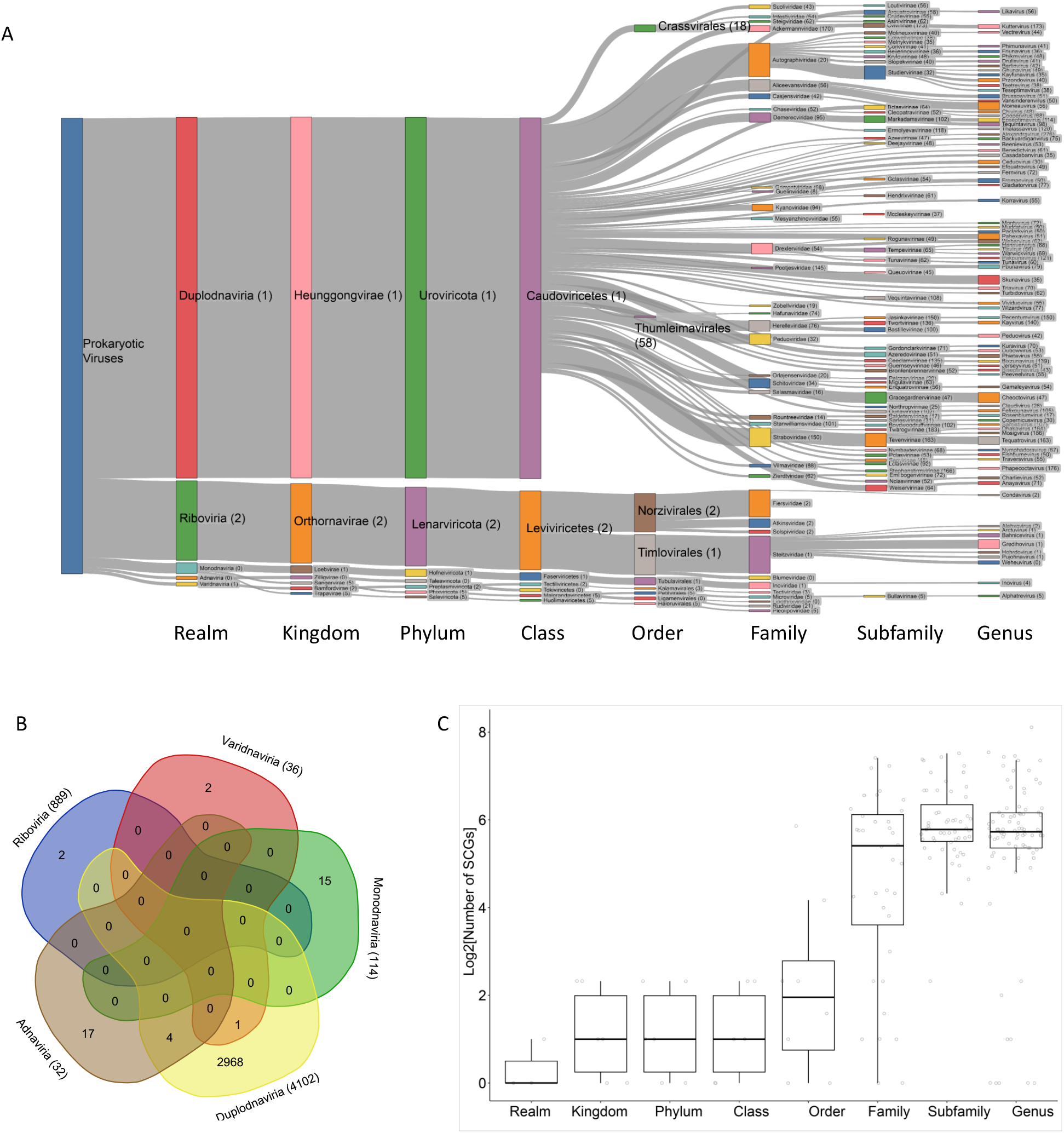
Single-copy genes (SCGs) of 203 taxa across prokaryotic viruses. (A) Number of single-copy marker genes of taxa. (B) Distribution of single-copy marker genes in viruses at realm level. The numbers in brackets indicate the total genomes analyzed for SCG identification within each realm. (C) Number of identified SCGs of taxa at each rank. This graph represents the number of SCGs identified in each taxon, depicted by individual circles. The median number of SCGs per taxon is shown by the horizontal line within each box. The upper and lower boundaries of the box indicate the upper and lower quartiles, respectively. The whiskers extend to 1.5 times the interquartile range, and data points beyond the whiskers are considered potential outliers.

There is a discernible pattern of fewer identified marker genes in selected taxa at higher taxonomic ranks (**Fig 2C**), as detailed in **Table S1**. All taxa possess no more than five genes at the order level or above, with only two at this level exhibiting more than five markers. In contrast, a significantly higher count of markers is observed in taxa at the family, subfamily, and genus ranks (**Fig 2C**). Among taxa at these lower ranks, 87.6% contain at least 10 markers, and 81.4% contain at least 30 markers. This inverse relationship between taxonomic rank and the number of identified single-copy marker genes can likely be attributed to the broader genetic variation encompassed within higher ranks. These ranks aggregate a wide array of genetically diverse groups, which dilutes the presence of common markers. Conversely, lower ranks, which represent more closely related entities, demonstrate greater genetic specificity, thus allowing for the identification of a higher number of unique markers. The scarcity of markers at higher taxonomic ranks challenges the resolution of deeper phylogenetic relationships, potentially necessitating supplementary methods for more accurate classification. The abundance of markers at lower ranks not only facilitates detailed taxonomic resolution within these groups but also enhances our understanding of their evolutionary relationships. For example, in the case of the family *Herelleviridae*, phylogenetic topologies based on 14 single marker genes effectively reveal evolutionary relationships at the subfamily and genus levels [36]. This underscores the utility of these markers in enriching our understanding of viral taxonomy and phylogeny.

### 3.2 High congruence with ICTV taxonomy at subfamily and genus ranks and with NCBI taxonomy at species rank

Based on the identified markers across the selected taxa, we constructed 203 phylogenetic trees, one for each of the 203 taxa analyzed. Despite this extensive effort, only the tree topologies for taxa at the family and subfamily levels — encompassing 36 families and 55 subfamilies — demonstrated high consistency with the ICTV classification system. This notable alignment underscores the robustness of the marker selection and tree construction methodologies at these taxonomic ranks. For a visual representation of this consistency, refer to the examples of phylogenetic tree topologies presented in **Fig 3**, and for further details, refer to **Table S3**. This specificity highlights the effectiveness of our approach in reflecting recognized taxonomic relationships at higher hierarchical levels, providing a strong foundation for reliable taxonomic classification within the framework established by the ICTV.

**Fig 3.**
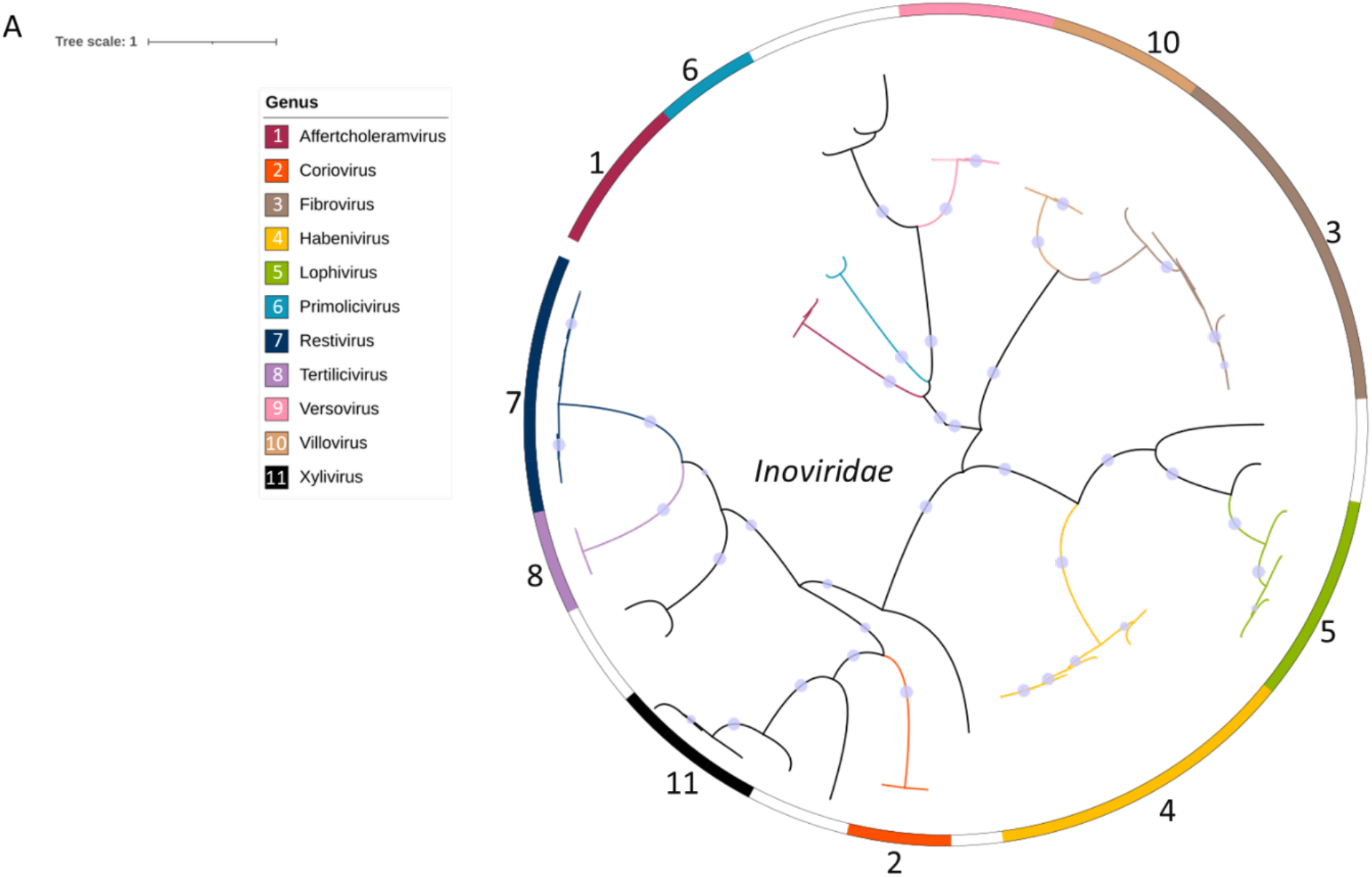
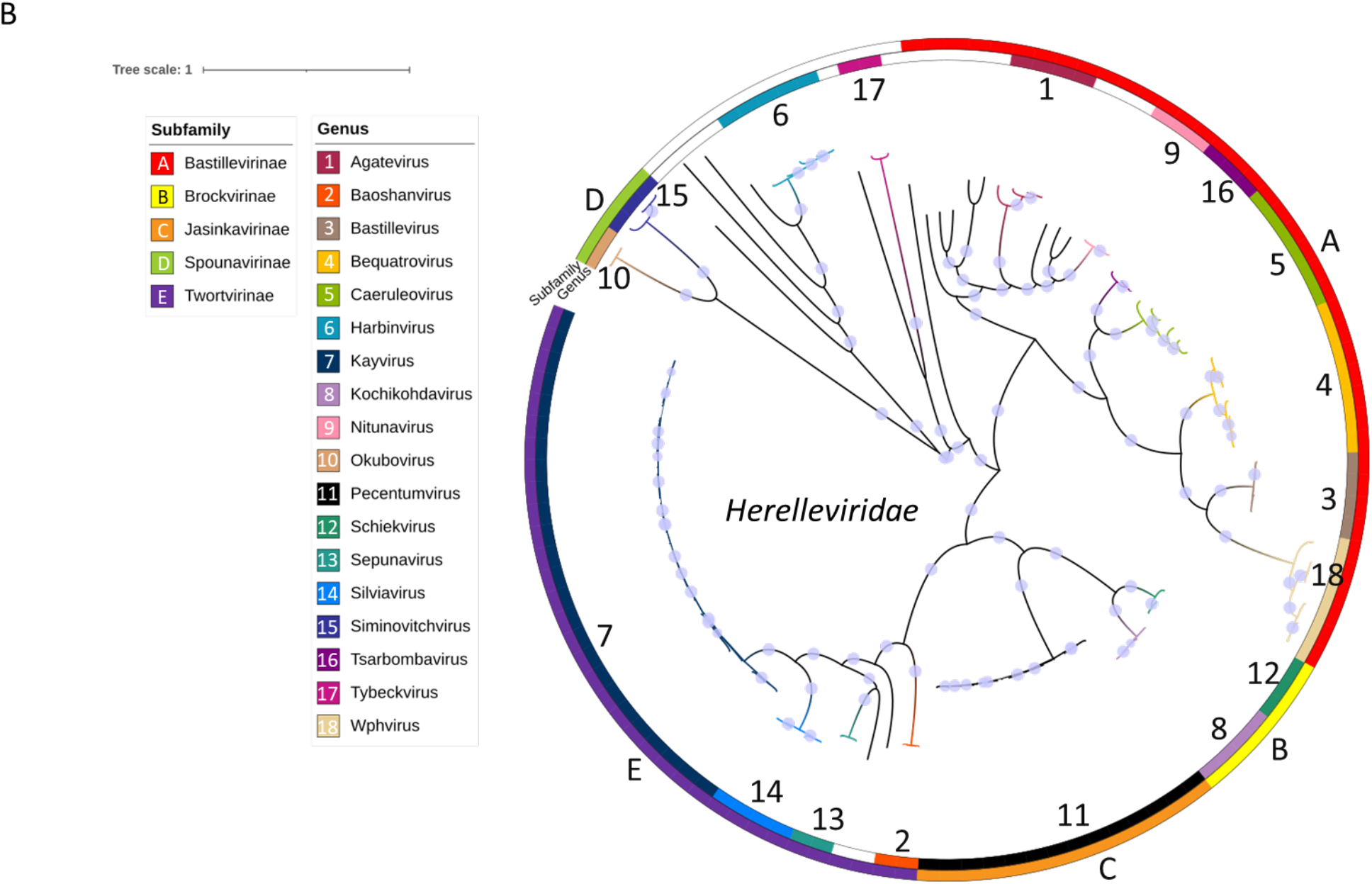
Representatives of phylogenetic tree topologies. (A) ssDNA representative: *Inoviridae* (100% consistent with ICTV taxonomy). (B) dsDNA representative: *Herelleviridae* (100% consistent with ICTV taxonomy). Solid blue circles indicate bootstrap values of 0.5 or higher. The tree is further detailed with color-coded rings; the outer ring represents viral subfamilies, and the inner ring denotes genera. Both subfamilies and genera within the *Inoviridae* and *Herelleviridae* families are identified as well-supported monophyletic groups, with bootstrap values of 0.75 or greater. This indicates a high level of congruence between the phylogenetic tree topologies and the ICTV classification, affirming the robustness of the phylogenetic structure in reflecting accurate taxonomic relationships.

For the 36 families analyzed, our results reveal substantial support for monophyly, with 85.9% of the taxa at the subfamily level (**Fig 4A**) and 84.3% at the genus level (**Fig 4B**) confirmed as monophyletic. This high level of monophyly is particularly pronounced among dsDNA viral families. For instance, families such as *Ackermannviridae, Herelleviridae*, and *Mesyanzhinovviridae* demonstrated exceptional congruence with ICTV taxonomy, with several achieving a perfect 100% alignment. Conversely, some families showed lower congruence values, such as *Aliceevansviridae* with only 33.3% congruence and *Peduoviridae* with 81.3%. The mixed-category *Pleolipoviridae* (dsDNA+ssDNA) exhibited a moderate congruence of 66.7%. In the realm of RNA viruses, congruence was notably variable, with *Fiersviridae* and *Steitzviridae* recording lower congruence values of 58.3% and 56.4%, respectively. These figures indicate classification challenges and discrepancies that could stem from the underlying phylogenetic methodologies or the existing ICTV taxonomic framework. The generally high congruence observed in many dsDNA viral families suggests that the employed phylogenetic method is reliable for this type of virus. However, the noted variability, especially among RNA viruses and specific dsDNA families, points to areas that could benefit from further refinement of the phylogenetic approach or necessitate updates to the ICTV taxonomic criteria to enhance accuracy and resolve existing discrepancies.

**Fig 4.**
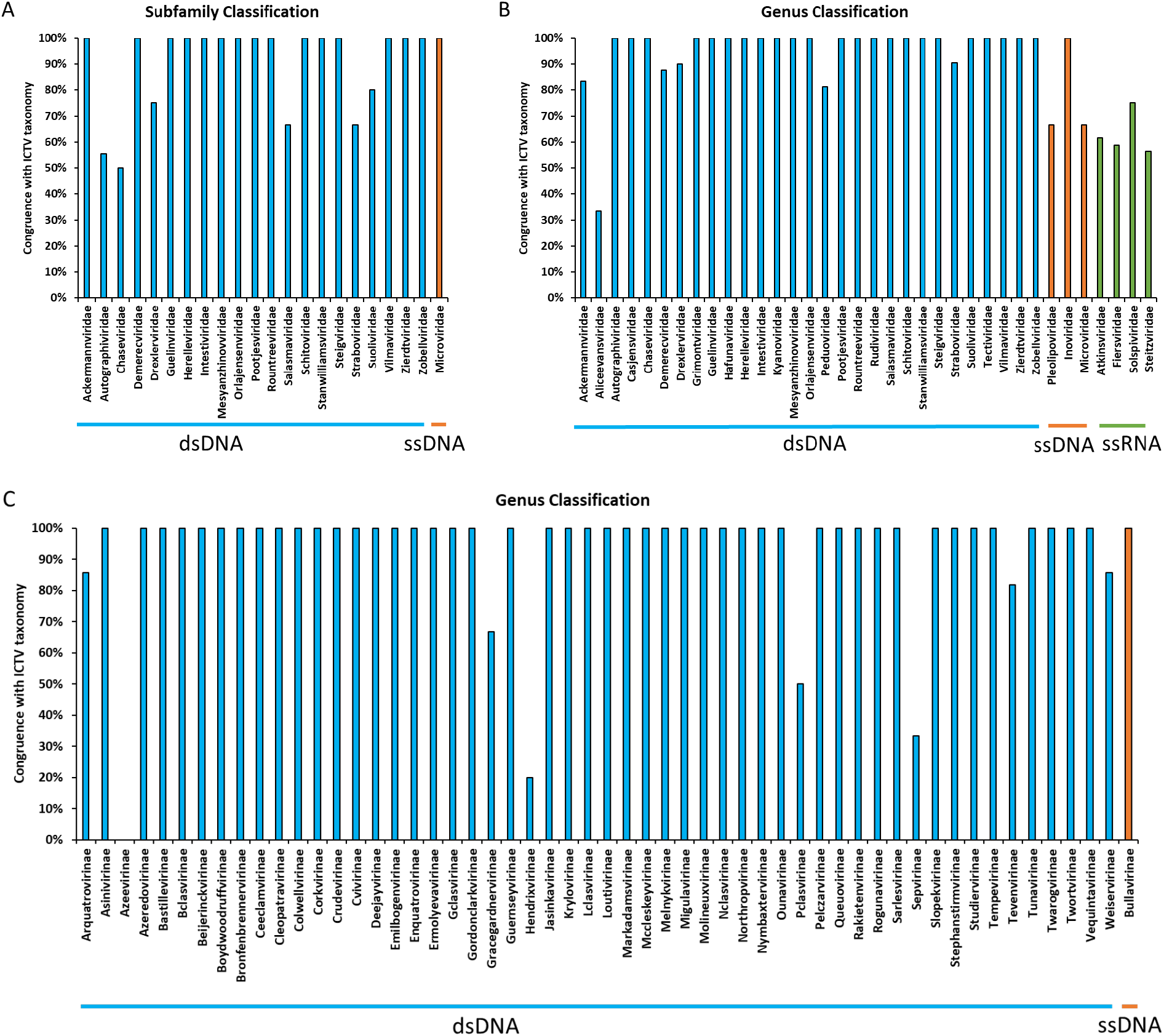
Congruence of phylogenetic tree topologies with ICTV subfamily and genus classification. Percentage bar charts in green color showing the consistency of tree topologies with ICTV subfamily (A) and genus (B) classification based on identified SCGs at family rank. Bars represent subfamily/genera with two or more genome representatives within a family (A and B). Percentage bar charts in blue show the consistency of tree topologies with ICTV genus (C) classification based on identified SCGs at subfamily rank. Bars represent genera with two or more genome representatives within a subfamily (C). Bar sizes represent the percentage of monophyletic subfamilies within a family or monophyletic genera within a family/subfamily.

For the 55 subfamilies analyzed, our results indicate strong monophyletic support at the genus level for 91.9% of the taxa (as shown in Fig **4C**). Notably, a perfect congruence rate of 100% was observed among the majority of dsDNA virus subfamilies, such as *Asinivirinae, Azedovirinae*, and *Bastillvirinae*. This high level of congruence underscores the consistency of the employed phylogenetic method with the ICTV taxonomy for these subfamilies, indicating a robust alignment. However, some subfamilies showed significantly lower congruence values, suggesting discrepancies in classification alignment with ICTV standards. Specifically, *Gorcegavirinae* exhibited a congruence of 66.7%, *Hendrixvirinae* had only 20%, *Pclasvirinae* had 50%, and *Sepvirinae* had 33.3%. These figures reflect a lesser degree of taxonomic alignment, which may prompt further investigations or necessitate revisions in the phylogenetic classification approach or the current ICTV taxonomy for these groups. The overall findings demonstrate a strong congruence for most dsDNA virus subfamilies, affirming the reliability and accuracy of the phylogenetic classification method utilized. However, the subfamilies with congruence values below 80% highlight areas of potential taxonomic uncertainty or ambiguity. These may stem from factors such as evolutionary divergence, lateral gene transfer, or other genetic complexities that are common in viral phylogenies, suggesting the need for continual refinement and adaptation of classification methodologies to accommodate evolving taxonomic insights.

Overall, the phylogenetic analysis of SCGs shows outstanding performance in classifying taxa at the subfamily and genus levels. However, when extending this approach to species-level classification, the congruence with ICTV taxonomy ranges between 67% and 75%. This indicates that relying solely on the phylogeny of SCGs may not be sufficiently effective for viral clustering at the species level. To enhance species classification accuracy, we adopted a combined approach that integrates the phylogeny of SCGs with ANI. For benchmarking, we selected GenBank genomes that contain species information and are affiliated with the specified 36 families or 55 subfamilies. Our method involved first placing these query genomes into reference trees to generate genera based on monophyly. Subsequently, we assigned species to these query genomes based on their ANI compared to reference genomes within the same genus. Since no other programs currently assign taxonomy at the species level, we exclusively utilized this combination approach to benchmark against NCBI taxonomy. Our results for the taxonomic assignment of 3,279 viral genomes (encompassing 1,920 species) from the 36 families showed that 1,772 out of 1,920 species (92.3%) aligned consistently with NCBI taxonomy. Similarly, the assignment of 2,703 viral genomes (1,450 species) from the 55 subfamilies demonstrated that 1,404 out of 1,450 species (96.8%) were consistent with NCBI taxonomy. These findings underscore the effectiveness of the combined approach of SCG phylogeny and ANI in achieving high congruence at the species level, thereby significantly advancing the precision and reliability of taxonomic classifications for viral genomes.

### 3.3 vClassifier was comparable to, or outperformed other tools

In comparison of subfamily/genus/species classification for the 36 viral families, we applied 2,564 genomes of 64 subfamilies belonging to the 36 families with reference phylogenetic trees to vConTACT3, VipTree, VirClust, and our approach (**Fig 5A**). For subfamily classifications, vClassifier showed the highest congruence (85.9%), followed by ViPTree (82.8%), VirClust (81.3%), and vConTACT3 (53.1%). For genus classification, ViPTree led with the highest congruence (88.2%), followed by vClassifier (84.3%), VirClust (68.5%), and vConTACT3 (48.9%). For species classification, vClassifier was the only tool that can assign species and showed significant congruence (92.3%). The vClassifier tool consistently exhibited the high congruence with the ICTV taxonomy at the subfamily and genus levels, and it was the only tool to demonstrate significant congruence at the species level. This suggests that vClassifier may be more accurate in capturing taxonomic nuances. The lack of congruence at the species level for vConTACT3, VirClust, and ViPTree could indicate limitations in their algorithms or databases, which might not be as comprehensive or recently updated as those used by vClassifier.

**Fig 5.**
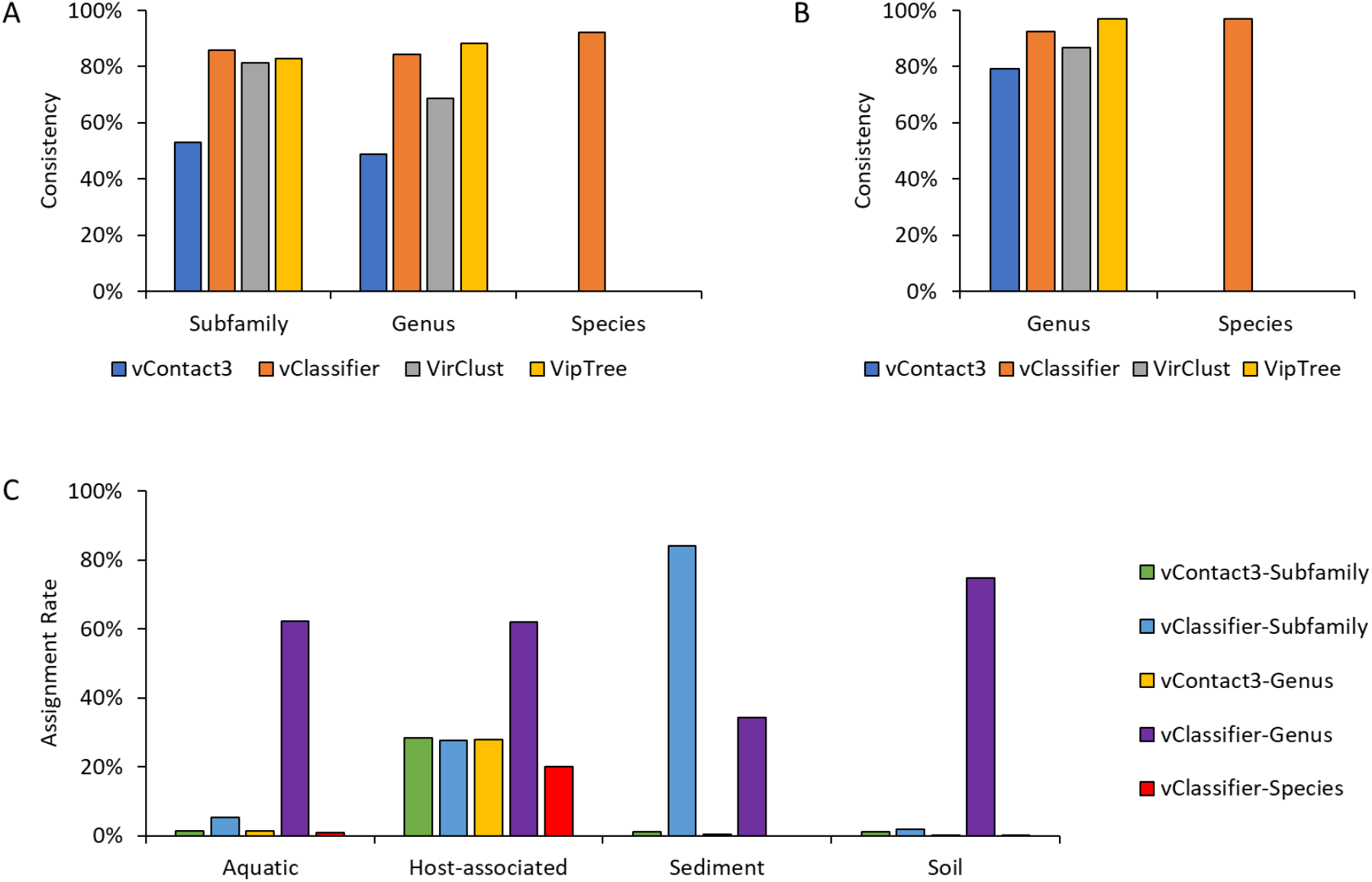
Comparative analysis of taxonomic consistency and assignment rates across different programs. Consistency of phylogenetic tree topologies with ICTV and NCBI classifications is shown for 36 families (A) and 55 subfamilies (B) that have ICTV classifications at the subfamily (A) and genus (A, B) levels as well as NCBI classifications at the species level (B). (C) Comparative taxonomic assignment rates of vClassifier and vConTACT3 across various environmental samples.

In our comparative study of genus and species classification for the 55 subfamilies, we analyzed 1,438 genomes across 172 genera using four different tools: vConTACT3, VipTree, VirClust, and our in-house developed vClassifier (illustrated in **Fig 5B**). At the genus level, VipTree exhibited the highest congruence at 97.1%. vClassifier also performed impressively, achieving a congruence of 92.4%. VirClust followed with a commendable congruence of 86.7%, while vConTACT3 demonstrated the lowest congruence among the tools at 79.1%. For species-level classification, vClassifier stood out as the sole tool offering this level of detailed classification, achieving a high congruence rate of 96.9%. The results at the genus level show a generally high level of congruence across all tools. While VipTree shows a slightly higher congruence than vClassifier at the genus level, it is important to note that VipTree does not incorporate bootstrap support in its analyses. The absence of this statistical support could lead to incorrect assignments, giving a false impression of higher congruence [12]. In contrast, the robust performance of vClassifier is achieved at both the genus and species levels and highlights its standardized analytical capabilities. The notable 96.8% congruence at the species level underscores vClassifier’s advanced utility for fine-scale taxonomic resolutions, which is essential for thorough viral characterization and research. The absence of species-level classification capabilities in the other tools could point to limitations in their current versions or a deliberate focus on broader taxonomic levels. This distinction emphasizes the unique position of vClassifier in providing detailed taxonomic insights, which can greatly enhance the precision of viral identification and classification studies.

### 3.4 Taxonomic assignment of viruses in diverse environments

In our comparative study between vClassifier and the well-known virus taxonomic assignment tool vConTACT3, we analyzed the assignment rates across various environmental samples (illustrated in **Fig 5C**). vClassifier demonstrated superior taxonomic assignment rates at the genus level for most environments. This was particularly notable in aquatic and soil samples, where it achieved assignment rates of 62.1% and 74.9%, respectively. vClassifier excelled at the subfamily level in sediment samples with an impressive 84.1% assignment rate, significantly outperforming other methods in this environment. In host-associated samples, both vConTACT3 and vClassifier showed relatively high assignment rates across all methods and levels. However, vClassifier had a higher rate at the genus level (62.1%) and maintained a decent rate at the species level (20.1%). Despite these strengths, vClassifier’s performance at the species level lagged compared to the genus level, with particularly low or no assignments noted in environments such as sediment and soil. Conversely, vConTACT3 generally showed lower assignment rates compared to vClassifier. Nonetheless, it provided some competitive performance at the host-associated subfamily level with an assignment rate of 28.47%. This comparison highlights vClassifier’s overall efficacy in assigning taxonomic classifications more accurately and effectively across various environmental contexts, especially at higher taxonomic levels, while also underscoring areas where improvements are needed, particularly in species-level classifications.

## 4 Discussion

Prokaryotic viruses display diverse evolutionary modes and rates, leading to notable differences in genome composition [37]. This variability impacts the conservation of markers across these viruses; notably, most single-gene markers are prevalent in fewer than 20% of viruses, which limits the phylogenetic signals they can provide [23]. This aligns with our observations that fewer single-copy marker genes are shared among taxa at higher ranks. While there are no universal single-copy genes across all viruses, specific sets of single-copy markers are frequently found within lower taxonomic ranks, such as families and subfamilies. Phylogenetic analyses based on concatenated markers at the family or subfamily level have consistently demonstrated strong monophyly of subfamilies and genera within our reference trees. These classifications correlate well with the standards set by ICTV, exhibiting over 80% consistency. However, the congruence between tree topologies and ICTV taxonomy drops below 80% when it comes to species classification. This discrepancy is likely due to the limited number of viral genomes that have been classified at the species level within the ICTV framework [38, 39], which may introduce biases in species classification assessments. To address this issue, we utilized an ANI approach for species classification, which has shown high consistency with the taxonomy provided by NCBI. The successful application of ANI, alongside phylogenetic analyses, suggests that combining these approaches offers a promising way to normalize rank assignments based solely on genomic data [19, 40]. This combination leverages robust phylogenetic frameworks and precise genomic comparisons, providing a comprehensive method for accurate and consistent viral taxonomy.

In this study, we developed vClassifier which leverages reference phylogenetic trees and ANI comparisons for determination of species-level taxonomy of viruses. vClassifier showed a high degree of consistency, exceeding 80%, with both ICTV and NCBI classifications. However, it’s crucial to acknowledge certain limitations of vClassifier regarding its classification capabilities. The tool is not designed for taxonomic classification of viral genomes at the family level or higher. Another limitation of vClassifier is that its capability for taxonomic classification only extends to the specific 36 families or 55 subfamilies recognized by the tool. While these constitute a large fraction of the families and sub-families on ICTV, vClassifier may not be able to generate useful outputs on novel viruses or viruses from other families outside of those represented in vClassifier’s database. To minimize these limitations, vClassifier will be updated regularly to ensure that its databases and taxonomy reflect the state of the art and to ensure users can fully exploit vClassifier’s capabilities to classify viruses at the subfamily, genus, and species ranks, thereby enhancing the overall scope and accuracy of viral taxonomy assignments.

## Acknowledgments

We thank members of the Anantharaman Laboratory for discussions and feedback on this software and manuscript

## Author contributions

KZ and KA conceived the project. KZ conducted bioinformatic analyses, statistical analyses, visualization of results, and content organization. KZ wrote and developed the code. KZ and KA wrote the manuscript draft. JK conducted data analyses and testing. All authors (KZ, KA, JK) reviewed the results, edited, and approved the manuscript.

## Code availability

Codes used in this project are available at the following GitHub repository: https://github.com/AnantharamanLab/vClassifier.

